# KIF18B is a cell-type specific regulator of spindle orientation in the epidermis

**DOI:** 10.1101/2021.06.03.446990

**Authors:** Rebecca S. Moreci, Terry Lechler

## Abstract

Proper spindle orientation is required for asymmetric cell division and the establishment of complex tissue architecture. In the developing epidermis, spindle orientation requires a conserved cortical protein complex of LGN/NuMA/dynein-dynactin. However, how microtubule dynamics are regulated to interact with this machinery and properly position the mitotic spindle is not fully understood. Furthermore, our understanding of the processes that link spindle orientation during asymmetric cell division to cell fate specification in distinct tissue contexts remains incomplete. We report a role for the microtubule catastrophe factor KIF18B in regulating microtubule dynamics to promote spindle orientation in keratinocytes. During mitosis, KIF18B accumulates at the cell cortex, colocalizing with the conserved spindle orientation machinery. *In vivo* we find that KIF18B is required for oriented cell divisions within the hair placode, the first stage of hair follicle morphogenesis, but is not essential in the interfollicular epidermis. Disrupting spindle orientation in the placode, using mutations in either KIF18B or NuMA, results in aberrant cell fate marker expression of hair follicle progenitor cells. These data functionally link spindle orientation to cell fate decisions during hair follicle morphogenesis. Taken together, our data demonstrate a role for regulated microtubule dynamics in spindle orientation in epidermal cells. This work also highlights the importance of spindle orientation during asymmetric cell division to dictate cell fate specification.

## INTRODUCTION

Spindle orientation is a fundamental biological process that contributes to asymmetric cell divisions and the formation of complex tissue architecture. Robust regulation of this process is essential for both the development and homeostasis of many tissues (Kulukian and Fuchs, 2013; Lu and Johnston, 2013; Poulson and Lechler, 2012). However, the mechanisms that regulate this process and connect spindle orientation to cell fate decisions in distinct tissue contexts remain incomplete.

In many cell types, spindle orientation/positioning requires a conserved cortical protein complex containing LGN/NuMA/dynein-dynactin that interacts with astral microtubules to properly orient the mitotic spindle. Previous work highlighted the necessity of dynein-dependent pulling for spindle oscillations, placement, and orientation. While targeting dynein to the cell cortex is sufficient to produce spindle oscillations, targeting both NuMA and dynein/dynactin to the cell cortex is required to produce asymmetric pulling forces to shift the mitotic spindle (Kotak et al., 2012; Okumura et al., 2018). In addition, there is a growing appreciation that precise control of microtubule dynamics is required for proper spindle orientation. Recent work in human osteosarcoma U2OS cells has shown that inhibition of the EB1-dependent plus-tip microtubule tracking protein GTSE1 is required to promote astral microtubule destabilization. As GTSE is inhibited, microtubule dynamic instability increases and enables astral microtubules to properly interact with the cell cortex and reposition the mitotic spindle during prometaphase (Bendre et al., 2016; Singh et al., 2021). Furthermore, the kinesin Kip3p, which acts as a microtubule depolymerase, promotes microtubule disassembly at the bud tip in budding yeast. Upon loss of Kip3p, microtubule length increased, resulting in mispositioning of the spindle pole body (Gupta et al., 2006). The mammalian homolog of Kip3p, Kif18b, has also been implicated in spindle positioning (McHugh et al., 2018). However, its role in spindle orientation and asymmetric cell divisions in intact tissue has not been examined.

Like Kip3p, KIF18B promotes microtubule catastrophe. In cultured mammalian cells, KIF18B is enriched at the plus tips of astral microtubules during mitosis, the microtubules that interact with the spindle orientation machinery (Supplemental Figure 1A) (Lee et al., 2010; Stout et al., 2011; Tanenbaum et al., 2011; van Heesbeen et al., 2017; Walczak et al., 2016). Intriguingly, KIF18B loss displays minimal effects on mitotic progression, therefore making mutants particularly viable resources for studying the role of microtubule dynamics during spindle positioning. Live imaging of HeLa cells demonstrated that loss of KIF18B results in increased spindle movements and displacement from the cell center during mitosis. Additionally, KIF18B knockout HeLa cells exhibited more spindle rotation in the z-plane (McHugh et al., 2018). These data support the idea that KIF18B is required for spindle positioning during cell division and that changes in microtubule dynamics can translate to defects in spindle positioning. However, the functional consequence of altered spindle positioning in this instance remains unclear. These models focused on roles of KIF18B in cells that exhibit non-polarized planar divisions; however, its role in regulating spindle orientation in polarized epithelia has yet to be examined.

**Figure 1.**
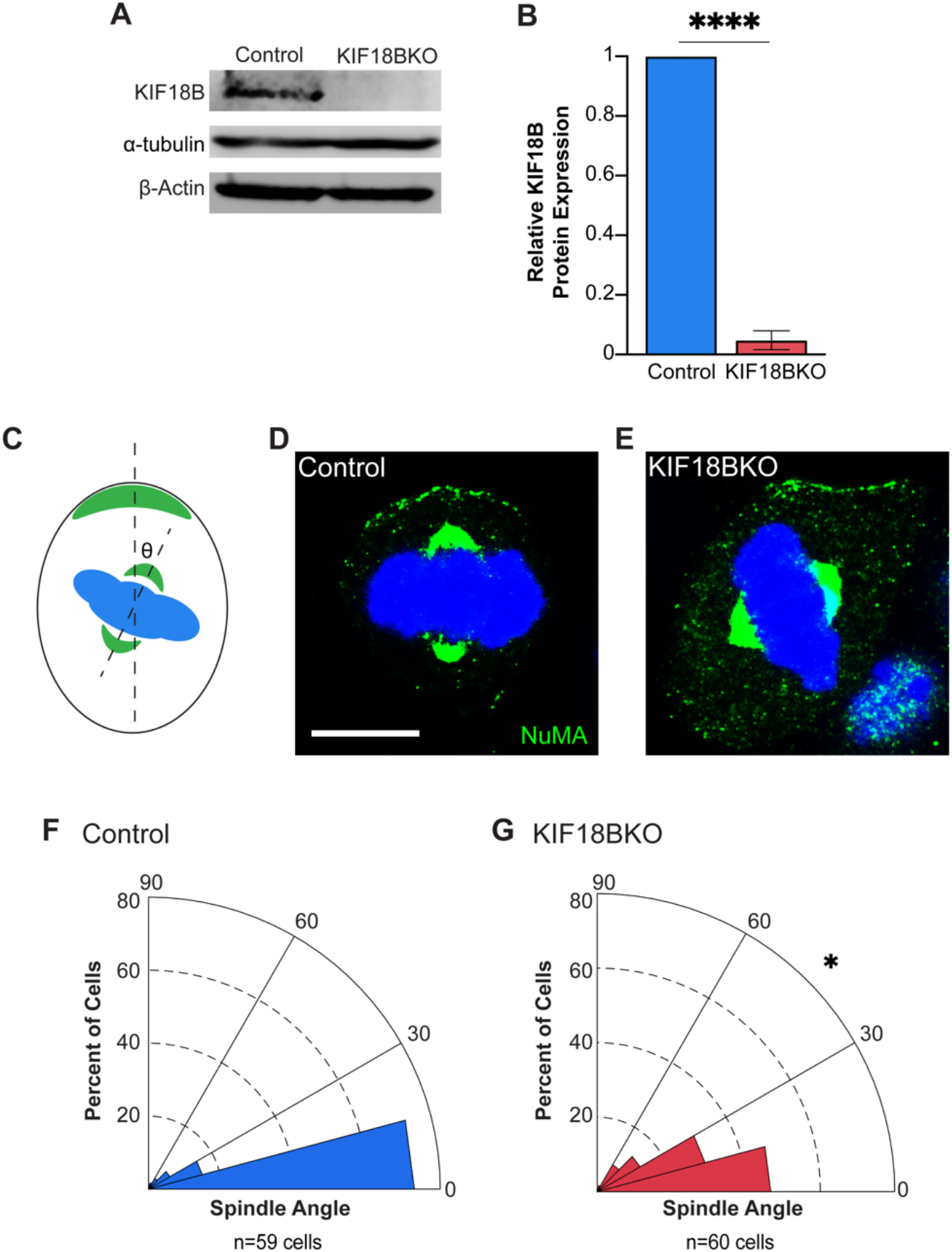
KIF18B is required for spindle orientation in cultured keratinocytes. (A) Western blot of proteins prepared from cell lysates for control and KIF18BKO keratinocyte cell lines. (B) Quantification of KIF18B protein levels in control and KIF18BKO (n=3, p < 0.0001), Student’s unpaired t-test). C) Diagram of spindle orientation measurement scheme with respect to cortical NuMA crescent (green). (D-E) Example images of spindle orientation with respect to NuMA (green) in control cells (aligned), KIF18BKO cells (misaligned) (F-G) Radial histograms of spindle orientation in control (F) and KIF18BKO (G) cells (p=0.0335, Kolmogorov-Smirnov Test). Scale bars = 10 μm.

The mammalian epidermis is an important model for studying the mechanism and function of spindle orientations *in vivo*. During development of the mammalian epidermis, oriented divisions of basal progenitors drive morphogenesis of the stratified epithelium, which is necessary for generating a barrier that shields against environmental insults, prevents water loss, and regulates ion exchange (Lechler and Fuchs, 2005; Poulson and Lechler, 2010, 2012; Smart, 1970). Defects in spindle orientation disrupt normal epidermal development and can lead to issues with barrier formation, differentiation defects, hair growth, and neonatal lethality, thus underscoring the importance of this process (Niessen et al., 2013; Seldin et al., 2016; Williams et al., 2011). The connection between spindle orientation and eventual cell fate in the epidermis has been demonstrated by both lineage tracing and live imaging (Box et al., 2019; Poulson and Lechler, 2012). Oriented divisions have also been observed in other cell types within the epidermis and its appendages, including placode cells during hair follicle morphogenesis and the transit amplifying cells of the hair matrix (Niessen et al., 2013; Ouspenskaia et al., 2016; Rompolas et al., 2012; Seldin et al., 2016). The hair placode is the first stage of hair follicle morphogenesis. In the hair placode, basal cells exclusively exhibit asymmetric divisions perpendicular to the underlying basement membrane. This division orientation has been proposed to be necessary for physically placing nascent suprabasal daughter cells away from dermal Wnt cues, enabling daughter cells to acquire a different fate (Ouspenskaia et al., 2016). However, physiological roles of orientated divisions in placode architecture and cell fate acquisition has not yet been shown.

Here, we examine the roles of KIF18B in both cultured keratinocytes and the developing epidermis. KIF18B mutants show an increase in both astral microtubule length and number as well as changes in microtubule dynamics. Furthermore, KIF18B colocalizes with spindle orientation machinery at the cell cortex. Using novel conditional loss of function mice, we demonstrate that KIF18B is required for spindle orientation during asymmetric cell division in hair placode compartments, but not in the interfollicular epidermis. Misorientation of placode spindles caused defects in cell fate decisions within the developing hair follicle. We propose that KIF18B regulates microtubule dynamics, enabling the spindle orientation machinery to properly interact with astral microtubules and align the mitotic spindle to promote robust cell fate decisions.

## RESULTS

### KIF18B is required for spindle alignment in cultured keratinocytes

To understand whether KIF18B is required for spindle alignment in keratinocytes, we first established control (Kif18b^floxed/floxed^) and KIF18B Knockout (KIF18BKO, Keratin-14Cre;Kif18b^floxed/floxed^) cell lines from neonatal mouse back skin. Knockout was established by use of a tissue-specific Keratin 14-Cre line to induced recombination in the basal layer of the developing epidermis (Vasioukhin et al., 1999). Recombination of the gene between the loxP sites removed a portion of the kinesin motor domain and induces a frameshift mutation (Supplemental Figure 1B). Western blotting demonstrated the loss of the KIF18B protein in KO cell lines (Figure 1A, B). To determine whether KIF18B was required for spindle orientation, we examined spindle alignment in control and KIF18BKO cultured keratinocytes. When keratinocytes cultured on laminin enter mitosis, they recruit spindle orientation machinery, such as NuMA, to a polarized cortical crescent (Seldin et al., 2016; Seldin et al., 2013). We defined the spindle alignment angle in each cell as the angle between a line connecting the two spindle poles and one that bisects the NuMA crescent on the cell membrane (Figure 1C). In control keratinocytes, the majority of cells divided with mitotic spindle angles within 15 degrees of the cortical NuMA crescent (Figure 1D, F). Loss of KIF18B resulted in an increased frequency of misaligned spindles in KIF18BKO cells compared to controls (Figure 1E, G). While loss of KIF18B does not cause complete randomization of spindle orientation like mutations in the cortical crescent machinery LGN, dynactin, or NuMA^ΔMTBD^, our data are consistent with KIF18B promoting the fidelity of spindle alignment (Seldin et al., 2016; Williams et al., 2011). The spindle orientation defect was also present in a second, independently isolated KIF18BKO cell line, confirming that the spindle orientation defect was not due to experimental technique (Supplemental Figure 1C-G). Notably, when KIF18B protein expression was restored with a full-length KIF18B tagged with a C-Terminal GFP, spindle alignment was rescued (Supplemental Figure 1C-G). Previous work determined that KIF18B’s regulation of astral microtubules was required for spindle centering and stabilization in planar cell divisions (McHugh et al., 2018). Our data establish a new role for KIF18B in promoting robust spindle orientation in keratinocytes to polarized intrinsic cues.

### KIF18B regulates microtubule length and number in keratinocytes

Despite the defects in spindle orientation found in KIF18BKO cells, the localization of the spindle orientation machinery components NuMA and dynein/dynactin (marked by p150^glued^, a subunit of dynactin) was normal (Supplemental Figure 2A-E). Compared to controls, KIF18BKO cells recruited NuMA and p150^glued^ at similar percentages as well as at similar levels, as marked by cortical fluorescence intensity (Supplemental Figure 2C-E), indicating that KIF18B more likely affects spindle orientation through a mechanism downstream of the localization of conserved cortical protein complex.

**Figure 2.**
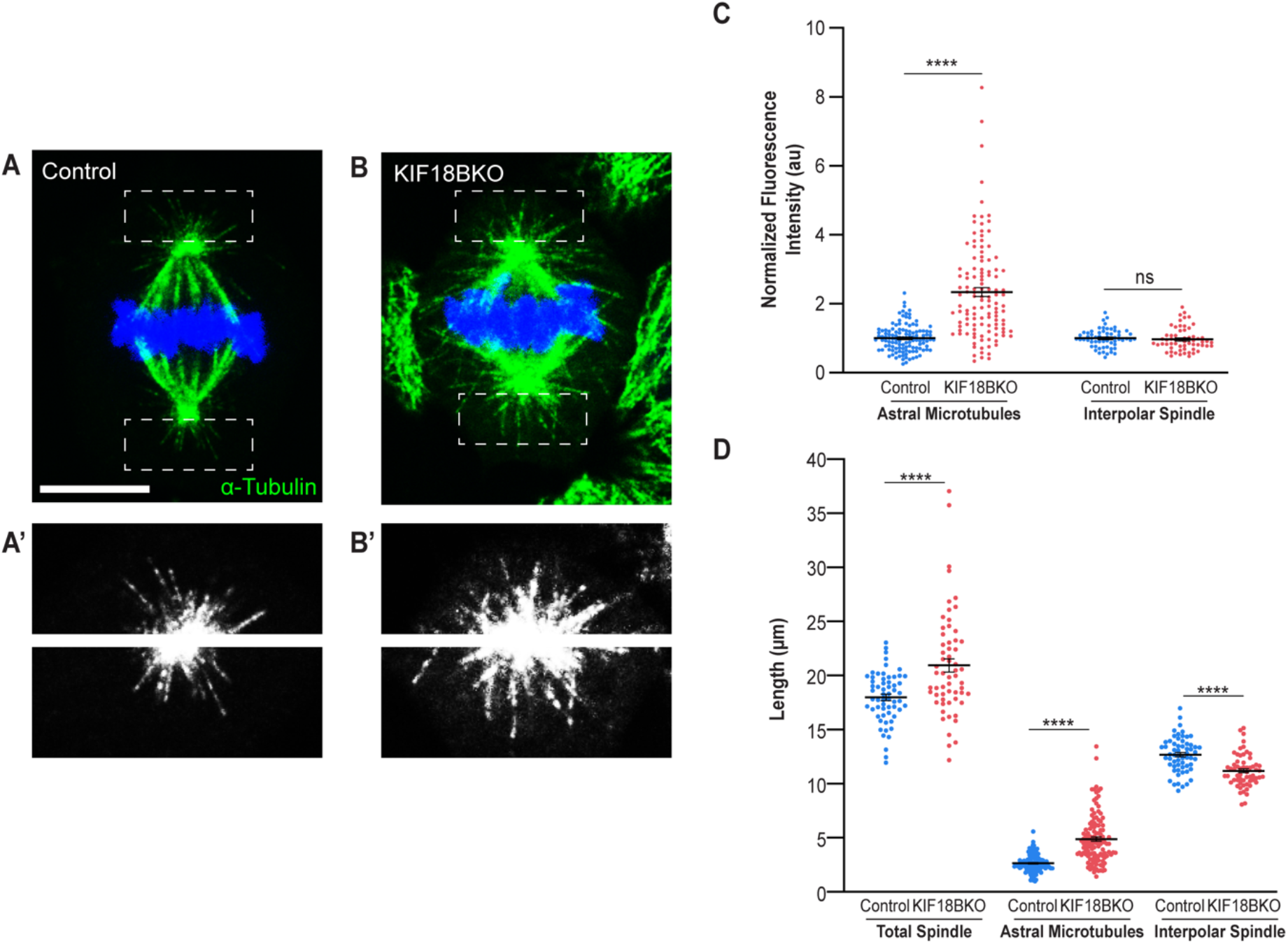
KIF18BKO alters spindle morphology in cultured keratinocytes. (A-B) Image of mitotic spindle during metaphase in KIF18BKO versus control keratinocytes. Alpha-tubulin (green) labels microtubules. (A’-B’) Gray-scale image of alpha-tubulin in control and KIF18BKO cells. (C) Quantification of microtubule fluorescence intensity at indicated spindle location, normalized to control cell average (n= 60 cells/condition, Astral Microtubules p<0.0001, Interpolar Spindle p=0.5614, Student’s unpaired t-test). (D) Quantification of spindle length at indicated location (n=60 cells/condition, Total Spindle p<0.0001, Astral Microtubules p<0.0001, Interpolar Spindle p<0.0001, Student’s unpaired t-test).

KIF18B promotes microtubule catastrophes *in vitro* (McHugh et al., 2018). Consistent with this, we found that loss of KIF18B produced a dramatic increase in both the length and number of astral microtubules compared to control mitotic spindles (Figure 2A-B’, Supplemental Figure 2F-H). Quantitation of microtubule fluorescence intensity demonstrated that astral microtubule number was greatly increased in the knockout versus the control, whereas fluorescence intensity was unchanged in the interpolar microtubules region, indicating that KIF18BKO predominantly affects astral microtubules (Figure 2B). The astral microtubule length of each spindle was increased between knockout and control cells while the length of the interpolar spindle was slightly shorter in KIF18BKO cells (Figure 2C, Supplemental Figure 2H). These data are in agreement with previous studies in other cell types (McHugh et al., 2018; Stout et al., 2011; Tanenbaum et al., 2011; Walczak et al., 2016). Both sides of the spindle were equally affected, as there was no difference in microtubule intensity or spindle length at either pole with respect to the cortical NuMA crescent (Supplemental Figure 2F-H) and KIF18B-GFP can partially rescue the spindle length (Supplemental Figure 2H).

### KIF18B regulates microtubule dynamics during mitosis

To better characterize the effects that loss of KIF18B has on astral microtubule dynamics, we next turned to live imaging. We isolated primary mouse keratinocytes from both control and KIF18BKO neonatal mouse back skin. These mice also expressed an EB1-GFP transgene to visualize polymerizing microtubules (Muroyama et al., 2016). KIF18BKO primary cells had more growing astral microtubules and longer astral microtubules compared to control cells. This became evident when examining the number of EB1 puncta in a single movie frame and standard deviation projections illustrating EB1 trajectories over time in KIF18BKO versus control cells (Figure 3A-C). Quantitation of the number of astral microtubules contacting the cell cortex revealed an almost 3X increase in KIF18BKO cells (Figure 3C). We then tracked EB1-GFP comets to gain insight into microtubule dynamics. KIF18B loss resulted in astral microtubules with increased growth distance (total μm traveled), growth lifetime (seconds visible in movie), and growth speed (μm traveled/second) (Figure 3D-F). These data are consistent with a decrease in microtubule catastrophes and demonstrate a requirement for KIF18B in regulating microtubule dynamics in keratinocytes.

**Figure 3.**
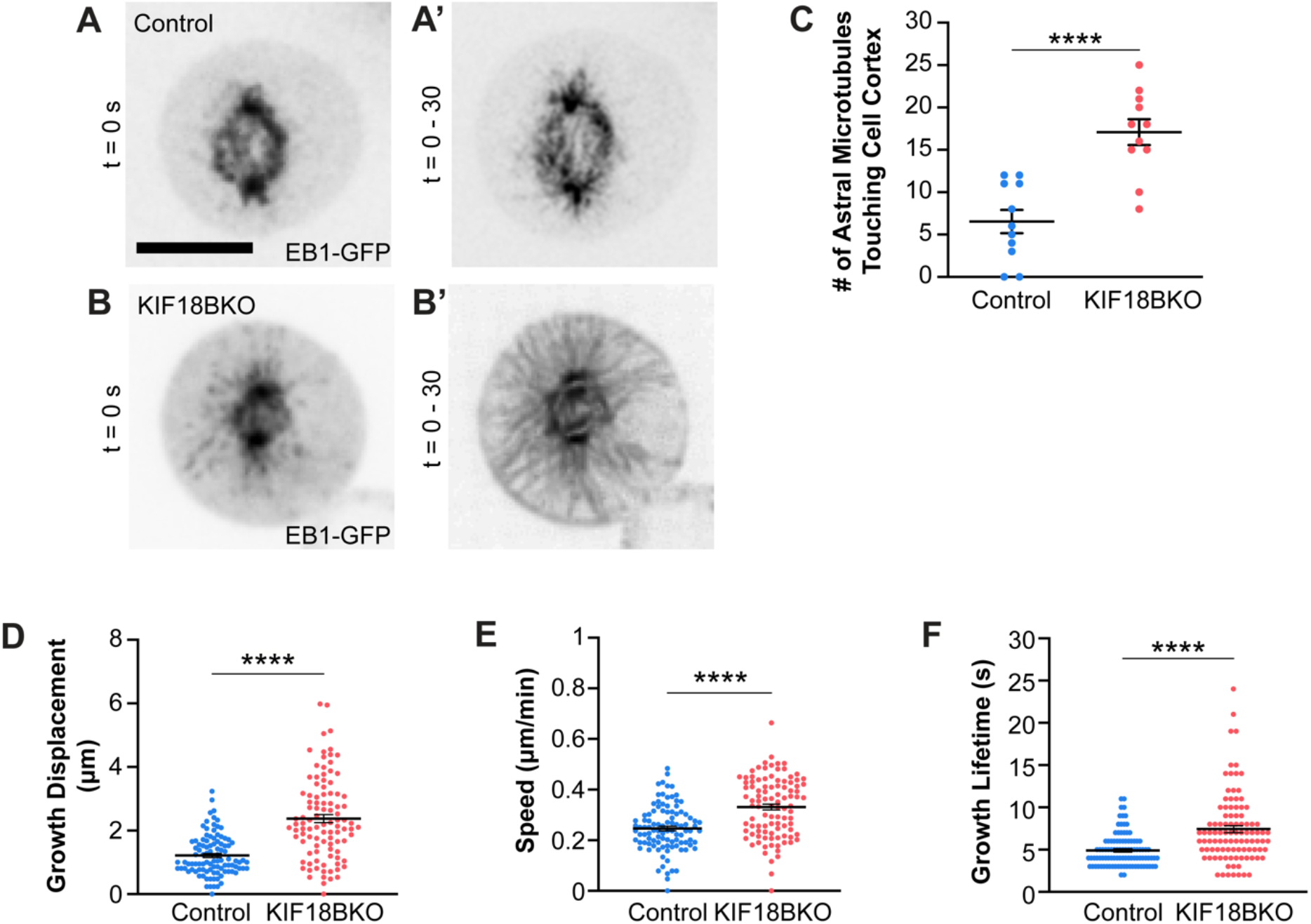
KIF18B regulates microtubule dynamics during mitosis. (A-B) Starting timepoint image (t=0) for EB1-GFP puncta from spinning disk movies in control and Kif18b KO cells. Movies are 30s, 1 frame/second. (A’-B’) Standard deviation projections of 30s movies. (C) Quantification of average number of astral microtubules touching cell cortex in control and KIF18BKO cells (n = 11 cells per condition, p <0.0001, Student’s unpaired t-test). (D) Growth displacement of EB1-GFP comets over 30s (n = 11 cells per condition, p<0.0001, Students unpaired t-test) (E) Speed of EB1-GFP comets over 30s time period (n = 11 cells per condition, p<0.0001, Student’s unpaired t-test). (F) Growth lifetime of EB1-GFP comets in control and KIF18BKO cells (n = 11 cells/condition, p<0.0001, Student’s unpaired t-test). Scale bar = 10 μm.

### KIF18B localizes to the cell cortex during mitosis in a NuMA-dependent manner

To better understand how KIF18B coordinates microtubule dynamics and spindle orientation, we turned to immunofluorescent staining. Immunolocalization of KIF18B in keratinocytes revealed a striking expression pattern; in addition to the astral microtubule tip localization that has been noted in other cell lines, we found that KIF18B localized to polarized crescents at the cell cortex during metaphase and notably colocalized with NuMA (Figure 4A-C). Both the spindle pole and cortical localization was absent in KIF18BKO cells, demonstrating the specificity of the antibody. Previous studies have not reported this localization pattern, which may reflect that those studies used cell lines that do not undergo polarization and spindle orientation to internal cues.

**Figure 4.**
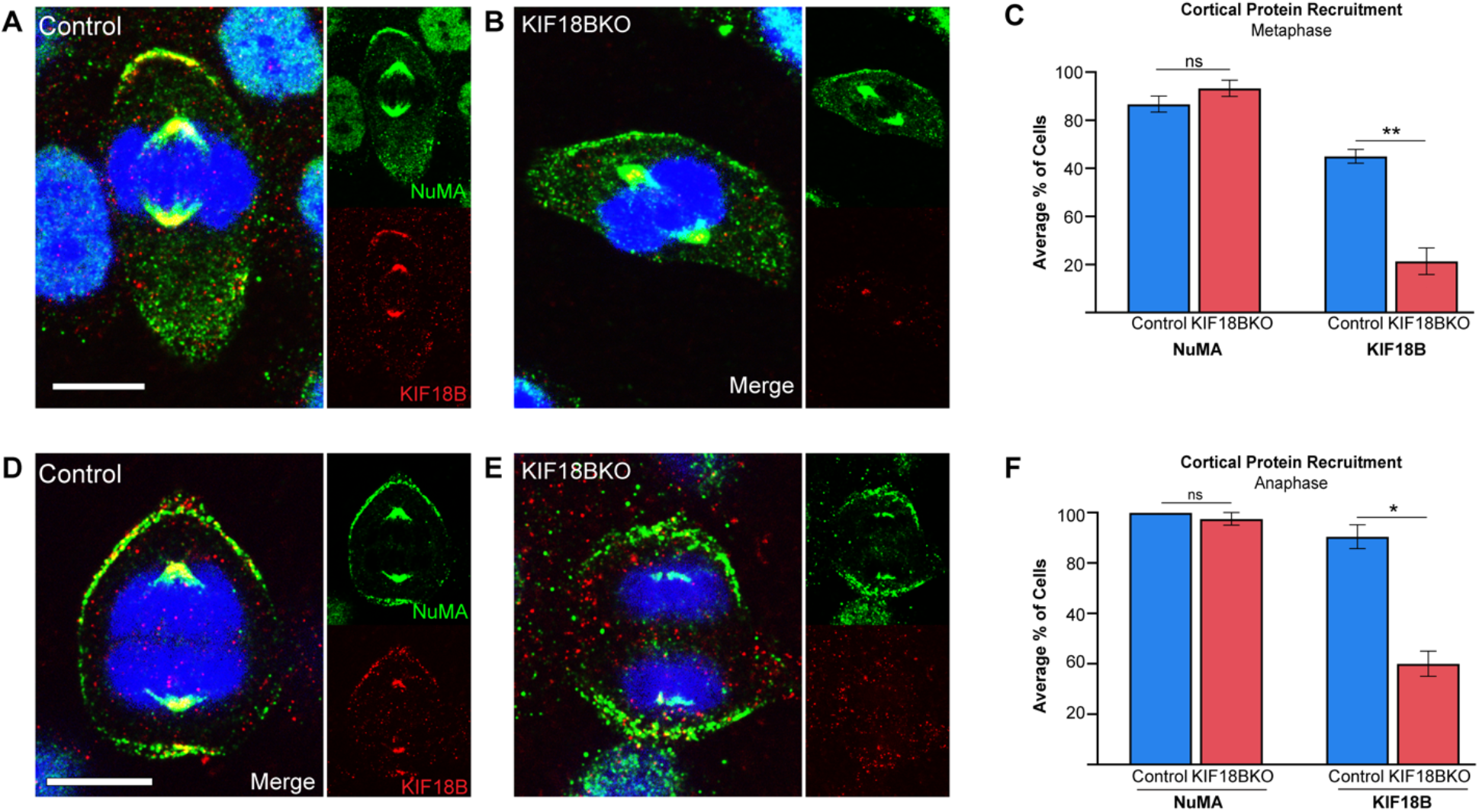
KIF18B accumulates at the cell cortex during mitosis. (A-B) Images of NuMA (green) and KIF18B (red) localization at the cell cortex in a crescent during metaphase. (C) Quantification of cortical protein recruitment in metaphase cells. (n=50 control cells, 48 KIF18BKO cells, NuMA p=0.2324, KIF18B p=0.0022, Student’s unpaired t-test). (D-E) Images of NuMA (green) and KIF18B (red) localizing cortically at both poles of the cell during anaphase. (F) Quantification of cortical protein recruitment in metaphase cells. (n= 40 cells/condition, NuMA p=0.4226, KIF18B p=0.0182, Student’s unpaired t-test). Scale bars = 10 μm.

Furthermore, KIF18B also colocalized with NuMA in anaphase. Previous work demonstrated that CDK1 phosphorylates NuMA during metaphase (Compton and Luo, 1995; Seldin et al., 2013). At anaphase entry, as CDK1 is inactivated, NuMA becomes dephosphorylated, and localization expands along the cell cortex to become bipolar. This same localization pattern was observed for KIF18B (Figure 4D-F).

As this cortical localization pattern has not been previously observed, we wanted to further probe how KIF18B is recruited to the cell cortex. NuMA cortical localization during metaphase requires direct binding to LGN (Seldin et al., 2013). We first utilized LGN knockdown (LGN KD) cells, which were previously created in the lab using validated shRNA constructs (Seldin et al., 2013; Williams et al., 2011). As expected, NuMA localization was absent at the cell cortex but was still present at the spindle poles during metaphase in LGN KD cells (Figure 5A-C). KIF18B exhibited the same expression pattern; upon loss of LGN, KIF18B no longer localized to the cell cortex during metaphase, but it was still present at the spindle poles (Figure 5A-C).

**Figure 5.**
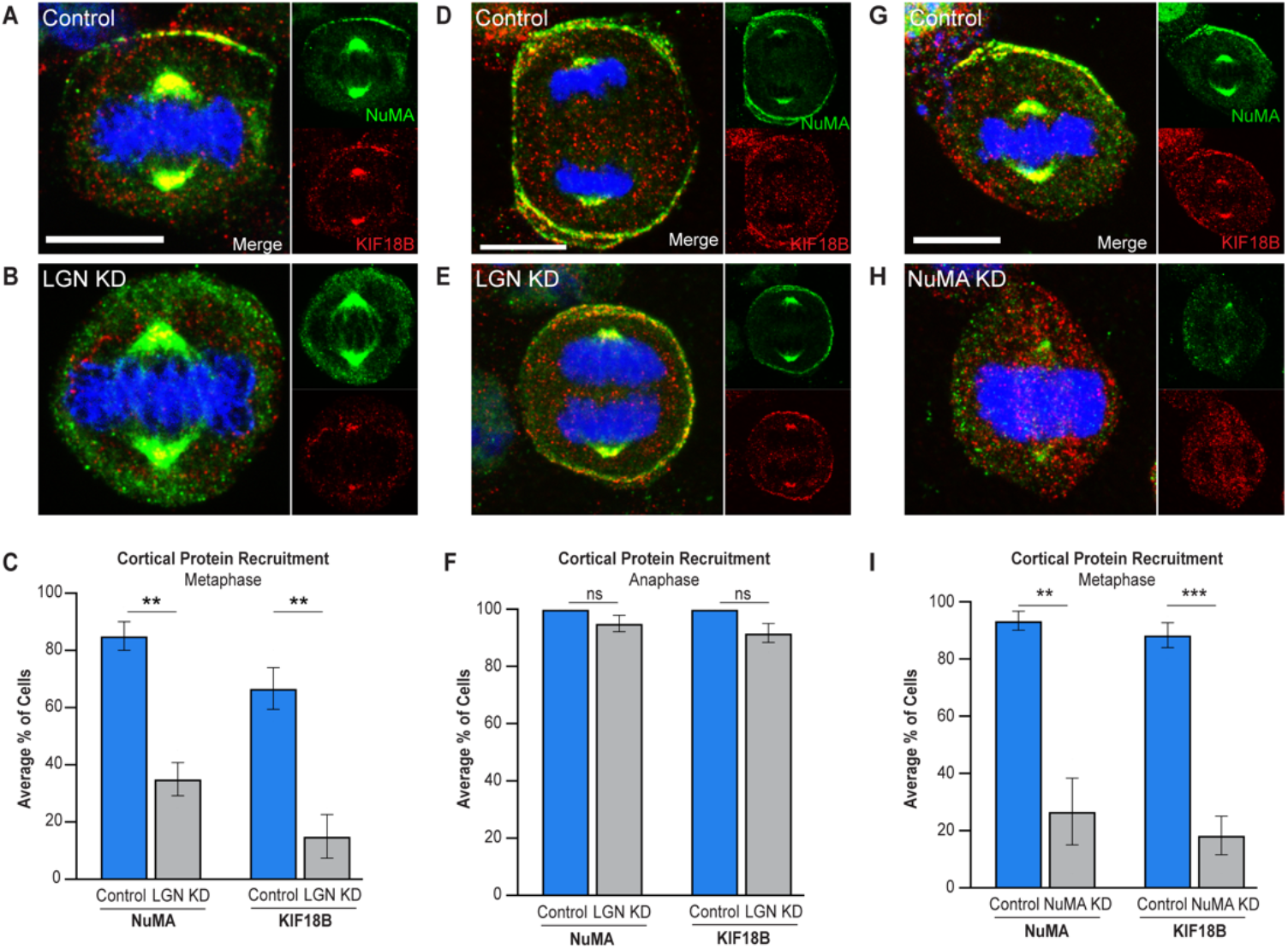
KIF18B localization requires spindle orientation machinery. (A-B) NuMA (green) and KIF18B (red) expression during metaphase in control and LGN knockdown cells (C) Graph of the percentage of metaphase cells with cortical NuMA and cortical KIF18B (n=60 cells/condition, NuMA p=0.0028, KIF18B p=0.008, Student’s t-test) (D-E) NuMA (green) and KIF18B (red) localization during anaphase in control and LGN knockdown cells. (F) Graph of the percentage of anaphase cells with bipolar cortical NuMA and cortical KIF18B (n=60 cells/condition, NuMA p = 0.1583, KIF18B p=0.0668, Student’s t-test) (G-H) NuMA (green) and KIF18B (red) localization during metaphase in NuMA knockdown vs. control cells (I) Graph of the percentage of metaphase cells with cortical NuMA and cortical KIF18B (n=60 cells/condition, NuMA p=0.0053, KIF18B p=0.0009, Student’s t-test). Scale bars = 10 μm.

As previously stated, NuMA’s localization is independent of LGN during anaphase, as CDK1 inactivation results in loss of NuMA phosphorylation, enabling it to bind directly to the cell membrane and accumulate in a bipolar fashion along the cell (Compton and Luo, 1995; Seldin et al., 2013). NuMA expression expanded along the cell cortex and was present at both cell poles in control and LGN KD cells (Figure 5D-F). Again, KIF18B localization mimicked that of NuMA, as it was present at both cell poles and spindle poles in LGN KD cells (Figure 5D-F). Consistent with the LGN-independent cortical localization of KIF18B and NuMA during anaphase, treatment of LGN KD cells with the CDK inhibitor Purvalanol A mimicked anaphase onset and promoted cortical localization of both NuMA and KIF18B during metaphase (Supplemental Figure 3A-C). These data suggest that like NuMA, KIF18B cortical localization is LGN-dependent during metaphase, but LGN-independent during anaphase.

We next tested whether the cortical KIF18B localization during mitosis was dependent on NuMA by utilizing previously established shRNA NuMA knockdown (NuMA KD) cell lines (Seldin et al., 2013; Williams et al., 2011). In these cells, NuMA staining was greatly diminished (Figure 5G-I). Further, KIF18B did not localize to either the cell cortex or the spindle poles, suggesting that KIF18B localization at both cell compartments is NuMA-dependent. To test if KIF18B cortical localization is dependent on microtubules, we treated cells with nocodazole to induce microtubule depolymerization. In DMSO-treated cells, control keratinocytes maintained cortical localization of both NuMA and KIF18B (Supplemental Figure 3D-F). In nocodazole-treated cells, spindle pole localization was disrupted, but cortical localization of both NuMA and KIF18B was preserved. Taken together, these data suggest that KIF18B localization at the cell cortex and spindle poles during mitosis requires NuMA. However, further investigation is needed to determine the function of cortical KIF18B, and if KIF18B binds directly to NuMA or forms a complex with additional proteins to promote its cortical localization.

### Spindle orientation is required for hair follicle morphogenesis

The requirement of cortical spindle orientation machinery for asymmetric cell divisions during epidermal development is well-documented (Seldin et al., 2016; Williams et al., 2011; Williams et al., 2014). Loss of LGN, NuMA, or dynein/dynactin results in a range of phenotypes, from barrier defects to neonatal lethality. Our data demonstrate that KIF18B is required to properly regulate microtubule dynamics to control spindle alignment in cultured keratinocytes. These *in vitro* observations led us to investigate if KIF18B also plays a role in regulating spindle orientation during epidermal development.

To understand the *in vivo* role of KIF18B in morphogenesis, we first examined KIF18B expression in the developing epidermis. During development, both the interfollicular epidermis and epidermal appendages, such as hair follicles, form in tandem. Placodes are small epidermal thickenings that mark the first stage of hair follicle morphogenesis. Interestingly, KIF18B expression was enriched in hair placodes at e16.5 during epidermal development (Figure 6A). Our KIF18B expression data is consistent with RNA-Seq datasets that also show placode enrichment compared to the interfollicular epidermis at the mRNA level (Rezza et al., 2016; Sennett et al., 2015). This is intriguing, as basal hair follicle cells have been shown to exclusively divide asymmetrically, with the mitotic spindle perpendicular to the underlying basement membrane (Ouspenskaia et al., 2016). These divisions were proposed to be functionally asymmetric, generating one slow-cycling basal cell and one suprabasal daughter cell marked by the transcription factor Sox9. Sox9-positive progenitors have been proposed to produce all lineages of the developing hair follicle, including the stem cell population within the bulge of the mature hair follicle (Nowak et al., 2008; Ouspenskaia et al., 2016). The functional role and mechanistic control of these placode asymmetric cell divisions remains unclear. Notably, previous work demonstrated that LGN, which is essential for spindle orientation of interfollicular cells, was present in placode cells though not required for the orientation of hair follicle progenitors (Byrd et al., 2016).

**Figure 6.**
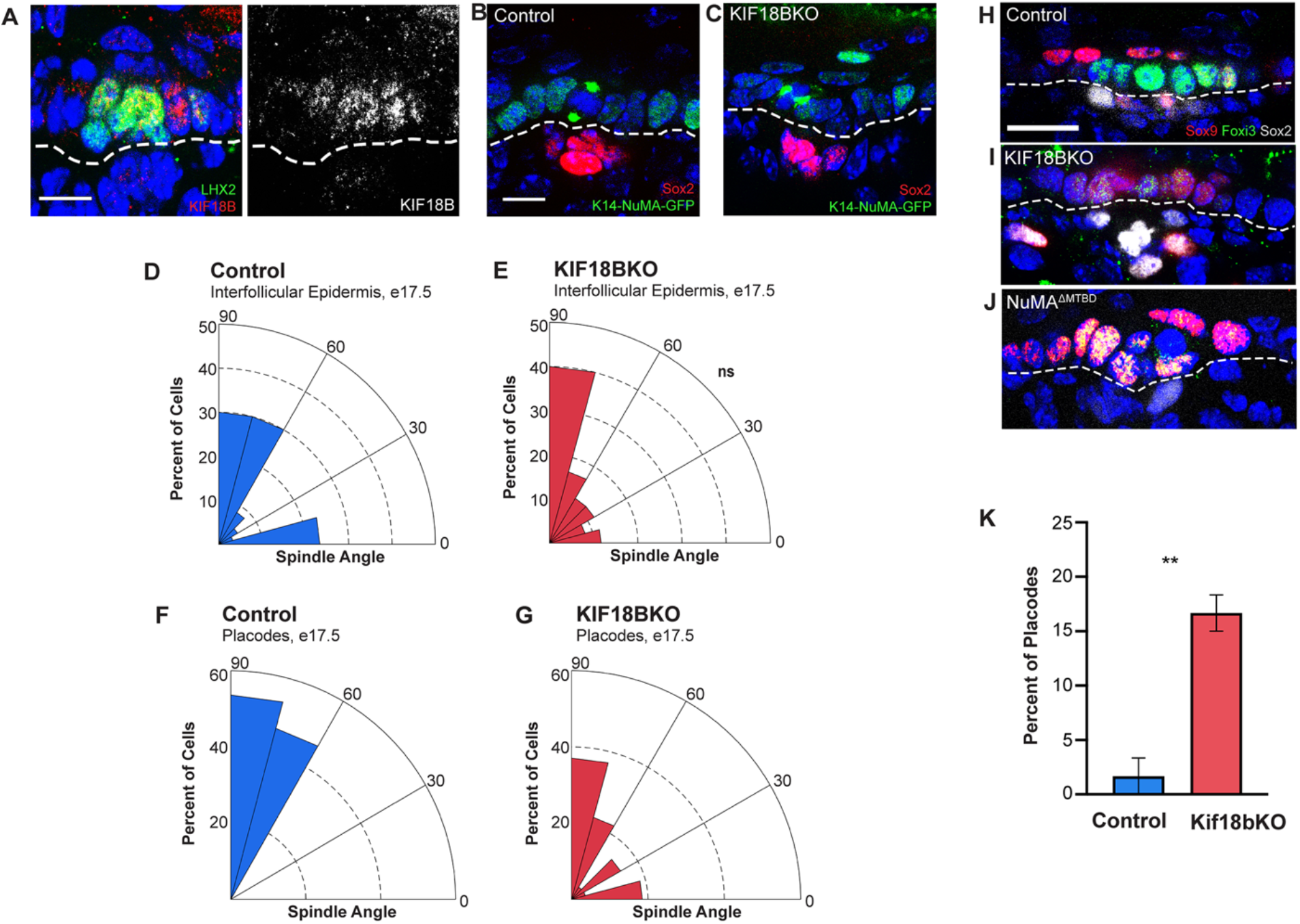
Spindle orientation is required for asymmetric cell division in hair placodes. (A) KIF18B (red) localization in hair placode at embryonic day 16.5. LHX2 (green) labels basal placode cells. Dotted line marks basement membrane. Scale bar is 10 μm. (B,C) Images of mitotic cells in control placode (B) and KIF18BKO placode (C). Sox2 (red) marks the dermal condensate while K14-NuMA-GFP (green) is used to measure spindle orientation with respect to basement membrane. Scale bar is 10 μm. (D-E) Radial histograms of spindle orientation in the back skin interfollicular epidermis of embryonic day 17.5 mice. (n=60 control cells, 60 KIF18BKO cells, p =0.1196, Kolmogorov-Smirnov test). (F-G) Radial histograms of spindle orientation in placode cells of e17.5 epidermis. (n=28 control cells, 27 KIF18BKO cells, p=0.0209, Kolmogorov-Smirnov test). (H) Control placode showing marker localization for progenitor cells (Sox9, red), basal cells (Foxi3, green) and dermal condensate (Sox2, grey). Scale bar is 20 μm. (I,J) Abnormal placodes from KIF18KO (I), and NuMA^ΔMTBD^ (J). (K) Quantitation of percent of mutant placodes in KIF18BKO compared to control. (n=60 placodes total per condition, 3 pairs of mice, p=0.0031, t test.

Next we assessed spindle orientation in both interfollicular and placode cells. We utilized the K14-Cre;Kif18b^fl/fl^ and also the K14-NuMA-GFP transgenic line to mark spindle poles and facilitate quantitation of division angles. We examined spindle orientation at embryonic day 17.5, a time point at which KIF18B is lost and at when the majority of spindles are perpendicular to the underlying basement membrane. *In vivo*, spindle orientation was measured by calculating the division angle of the two spindle poles (marked with NuMA-GFP) with respect to the basement membrane. Spindle orientation within the interfollicular epidermis was not statistically significantly different between KIF18B KO and control; the majority of divisions remained asymmetric with spindle angles between 60-90 degrees (Figure 6D,E). This is consistent with the low levels of KIF18B expression in these cells. In contrast, when we examined spindle orientation in the hair placodes (using Sox2 to labeling to identify these structures), we observed a clear role for KIF18B (Figure 6B,C,F,G). In control placodes, the majority of basal cells divided perpendicularly, as previously reported with division angles between 60-90 degrees (Figure 6F) (Byrd et al., 2016; Ouspenskaia et al., 2016). Strikingly, KIF18BKO placodes contained basal cells dividing parallel to the basement membrane and across the spectrum of 0-90 degrees (Figure 6G). Division angle did not correlate with position within the placode, as parallel-dividing cells were present in both the center and at the edges of placodes. This specific role for KIF18B in placode spindle orientation allows us to probe the functional role of these oriented divisions for the first time.

Previous studies proposed that division of Wnt^high^ basal cells away from the basement membrane is sufficient to drive changes in cell fate, establishing Sox9^positive^/Wnt^low^ expression within suprabasal placode cells (Ouspenskaia et al., 2016). This model assumed that cell fate specification is driven by external factors, and that division into a Wnt^low^ environment instructs cell fate decisions. We tested this model using our conditional KO mouse. We first examined cell fate markers within control placodes. As expected, control placodes exhibited the stereotypical staining pattern, with Sox2 marking dermal condensates and the transcription factor Foxi3 marking basal placode cells. Sox9 nuclei were present within the suprabasal layer of the placode (Figure 6H). Surprisingly, when we stained for these same markers in KIF18BKO mice, we observed Sox9 expression in the basal layer of the hair placode, an area from which it is normally absent (Figure 6I). To quantify this phenotype, we classified placodes as abnormal if they contained three or more Sox9^positive^ cells within the basal layer, as border placode cells display occasional Sox9 expression. In KIF18BKO mice 15-20% of placodes were mutant (Figure 6K). To understand if this phenotype was specific to KIF18B mutants or a more general asymmetric cell division phenotype, we examined placodes in another asymmetric cell division mutant. We previously showed that a mutant of NuMA lacking its microtubule-binding domain (NuMA^ΔMTBD^) had defects in interfollicular spindle orientation (Seldin et al., 2016). NuMA^ΔMTBD^ mutants demonstrated the same placode phenotype (40% of placodes abnormal, n=20), with Sox9 expression expanding into the basal layer of the placode (Figure 6J). Taken together, these data suggest that disruption of spindle orientation and asymmetric cell division within hair placodes leads to changes in the architecture/organization of these structures, consistent with defects in asymmetric cell divisions and cell fate specification.

## DISCUSSION

In this study, we characterized the role of the kinesin KIF18B in spindle orientation in keratinocytes and during skin and hair follicle morphogenesis. Loss of KIF18B increased astral microtubule length, amount of astral microtubule cortical contacts, and led to defects in spindle orientation in cultured keratinocytes. Intriguingly, KIF18B is specifically required for spindle orientation and cell fate specification in hair placodes but dispensable in the interfollicular epidermis, highlighting the requirement of distinct cortical machinery in different tissue contexts.

Our work demonstrates KIF18B’s roles in controlling microtubule dynamics and spindle morphology in keratinocytes. Consistent with previous studies, disrupting KIF18B affected the morphology and dynamics of astral microtubules in epidermal keratinocytes (Lee et al., 2010; McHugh et al., 2018; Stout et al., 2011; Tanenbaum et al., 2011; Walczak et al., 2016). KIF18BKO keratinocytes showed an increase in length and number of astral microtubules while EB1-GFP tracking revealed that astral microtubules contact the cell cortex more often in KIF18BKO cells than in control cells. EB1 puncta also exhibited larger growth displacements and growth lifetimes in KIF18BKO cells compared to controls. Taken together with the spindle orientation results, our data suggest that KIF18B promotes proper spindle orientation by modulating microtubule dynamics. Microtubule dynamics are also known to contribute to spindle positioning in other contexts, such as the one-cell stage of *C. elegans* development and budding yeast (Gupta et al. 2006;Nguyen-Ngoc et al., 2007).

In metazoans, the conserved cortical spindle orientation machinery generates pulling forces on astral microtubules to properly rotate the spindle into place (Lu and Johnston, 2013; Okumura et al., 2018). Our data demonstrated that KIF18B co-localizes with the cortical spindle orientation machinery in keratinocytes. KIF18B localization mimics that of NuMA, requiring LGN for cortical localization during metaphase but not anaphase. KIF18B required NuMA to localize at both the spindle poles and the cell cortex during metaphase and anaphase. Whether KIF18B and NuMA bind directly to one another or form a complex with other proteins will require further investigation. However, this raises questions about whether coordination of microtubule dynamics locally at the cell cortex, globally throughout the cell, or both are required for proper spindle orientation. Future experiments aimed at specifically disrupting KIF18B’s cortical localization should address this. Given that NuMA’s microtubule binding domain and interactions with astral microtubules are required for robust spindle orientation, it is intriguing to speculate that KIF18B’s depolymerizing activity at plus ends of astral microtubules can be coopted by cortical machinery to guide spindle orientation and movement (Seldin et al., 2016).

Previous work identified KIF18B as a regulator of spindle positioning and centering in HeLa cells (McHugh et al., 2018). That work suggested that KIF18B is necessary for the planar alignment of the mitotic spindle with the underlying substrate. However, until now, KIF18B’s role in regulating spindle orientation in cells that undergo polarized, asymmetric cell divisions had not been tested. We demonstrated that KIF18B is required for spindle orientation in cultured keratinocytes. However, we found no role for KIF18B in the planar or perpendicular divisions within the interfollicular epidermis. This is surprising for two reasons. First, the work in HeLa cells predicted that KIF18B would be requires for planar spindle orientation of epithelial cells. Second, to our knowledge, KIF18B is the first example of a protein that is required for spindle orientation in cultured keratinocytes, but not in interfollicular epidermal cells *in vivo*.

In contrast, KIF18B has a clear role in spindle orientation in hair placodes during hair follicle morphogenesis. Previous research had observed oriented divisions in the placode, but the function and regulation of these divisions was not directly tested (Ouspenskaia et al., 2016; Williams et al., 2011). Interestingly, LGN is required for perpendicular spindles in the interfollicular epidermis but dispensable in hair placodes, while NuMA is required for both (Byrd et al., 2016). These data emphasize the unique requirements for spindle orientation machinery amongst different sub-compartments of the epidermis. Whether these distinctions reflect fundamental differences in microtubule dynamics or distinct regulators of microtubule dynamics between different pools of epidermal stem cells is a fascinating question that requires further attention.

The long-term effects of this mutant placode phenotype remain unclear and require further investigation. Hair follicle morphogenesis occurs in waves, with the first wave beginning around embryonic day 12 or 13 (Saxena et al., 2019). The K14-Cre promoter used here begins to turn on around e14.5, and proteins are lost by e15.5-16.5. Therefore, many hair placodes form before recombination, preventing us from exploring how KIF18B loss, and particularly disruptions in placode spindle orientation, impacts overall hair follicle morphogenesis. While we were generating the KIF18B^fl/fl^ mouse line, we noted that KIF18B mice containing the Neo cassette acted as hypomorphs and displayed the same hair placode phenotype (KIF18B^Neo^), even at earlier developmental timepoints (Supplemental Figure 4A-C). Scenarios in which intact Neo cassettes disrupt regular gene expression have been well-documented (Clark et al., 2020; Meier et al., 2010; West et al., 2016). We were only been able to collect a small number of these samples due to mid-embryonic lethality. However, these hypomorphs exhibited a decrease in hair follicles at embryonic day 18.5 (Supplemental Figure 4D-F). This line does not allow us to determine whether this effect is cell autonomous, however, and future work will be needed to determine the fates of placodes with disrupted patterning.

A current model suggests that oriented divisions out of a Wnt^high^ basal signaling hub are sufficient to drive cell fate changes in the placode (Ouspenskaia et al., 2016). In support of this, recent work identified asymmetric secretion of both Wnt repressors and activators within the placode (Matos et al., 2020). However, analysis of KIF18BKO placodes suggests novel modes of intrinsic control of cell fate in addition to extrinsic cues. In the mutant, Sox9 progenitor cells were present within the basal placode layer, adjacent to dermal Wnt signals. If cell fate control were under extrinsic control alone, these cells should have the correct fate. Furthermore, we were able to recapitulate expansion of Sox9 expression into the basal layer when treating mice with Porcupine inhibitor LGK974 (Supplemental Figure 4G-H). Together, these data are consistent with a model in which Wnt regulators, whether repressors or activators, are asymmetrically inherited during oriented cell divisions to dictate cell fate specification. An alternative explanation is that KIF18B directly affects Wnt signaling. While we cannot rule this out, the fact that both NuMA and Kif18b mutants disrupt spindle orientation and cell fate acquisition suggests a more direct role for asymmetric cell divisions. This also highlights the importance of identifying asymmetrically inherited factors that influence these fate decisions. Collectively, our work demonstrates a functional role for regulated microtubule dynamics during asymmetric cell division and the importance of oriented divisions during epidermal development.

## MATERIALS AND METHODS

### Mice

All animal work was approved by Duke University’s Institutional Animal Care and Use Committee. Mice were maintained in a clean, barrier facility with 12 hr light/dark cycles. Both male and female mice were used in this study. Mice were genotyped by PCR to confirm strain before analysis. Mouse strains used in this study were: Kif18b^fl/fl^ (generated by Duke Transgenic and Knockout Mouse Shared Resource), Keratin 14-Cre (Vasioukhin et al., 1999); K14-EB1-GFP (Muroyama et al., 2016); Keratin 14-NuMA-GFP (Poulson and Lechler, 2010), NuMA1-ΔMTBD (Silk et al., 2009).

### Cell Culture

Keratinocytes were maintained at 37 °C, 7.5 % CO_2_ and grown in E low calcium medium. Cells were incubated with additional 0.5 mM calcium for 4 hours before experimental analysis. To produce primary keratinocytes, back skin was removed from mice (p0-p3) and placed in 1:1 Dispase II (Hoffman-La Roche, Basel, Switzerland):1X phosphate buffered saline (PBS) solution at 4 °C overnight. Epidermis was then separated from the dermis and placed in a 1:1 mixture of versene/trypsin-EDTA (0.25%) for 12 minutes. Following incubation, cells were resuspended in medium, filtered (70 μm cell strainer), pelleted, and resuspended in medium before plating. For immediate experimental use, cells were plated on 3.5 mm glass-bottomed TC plates coated with laminin. Medium for primary cultures was supplemented with 0.5 mM calcium and Dual SMAD inhibitors (Mou et al., 2016). To create stable lines, cells were initially plated with 0.5 mM calcium onto fibroblast feeders. With each passage, calcium content was decreased by 0.1 mM until cells were able to be weaned off calcium and feeders. KIF18B control and KO cell lines were additionally grown with dual SMAD inhibitors.

LGN Knockdown and NuMA knockdown cells were previously made in the lab using siRNA constructs (Seldin et al., 2013). Early passages (p0-p3) of both stable lines were used. Culture was supplemented with puromycin to maintain knockdown.

Phoenix cells were passaged at 37 °C, 7.5 % CO_2_ grown in DMEM/10% FBS but switched to 32 °C to produce retrovirus.

### Drug and Chemical Information

| Drug | Supplier | Catalog # | Use | Working Concentration | Treatment Time |
| --- | --- | --- | --- | --- | --- |
| A 83-01 | Sigma-Aldrich | SML0788 | Dual SMAD Inhibitor Cocktail | 1 µM | Continuous |
| CHIR99021 | Sigma-Aldrich | SML1046 | Dual SMAD Inhibitor Cocktail | 1 µM | Continuous |
| DMH-1 | Sigma-Aldrich | D8946 | Dual SMAD Inhibitor Cocktail | 1 µM | Continuous |
| LGK 974 | Caymen Chemical | 14072 | Wnt Inhibitor, IP Injections | 3 mg/kg | e15.75, e16.25 |
| Nocodazole | Sigma-Aldrich | M1404 | Depolymerize microtubules | 10 µM | 10 min |
| Puromycin | Amresco | J593-25MG | Selection for cell lines | 2 µg/mL | Continuous |
| Purvalanol A | Tocris | 1580 | CDK Inhibitor, Cell Culture | 100 µM | 3 min |
| Y-27632 | Selleck Chemicals | S1049 | Dual SMAD Inhibitor Cocktail | 5 µM | Continuous |

### Production of KIF18B-GFP Construct and Cell Line

All PCR for cloning purposes was performed using Verifire DNA Polymerase. GFP was amplified from the pEGFP-C1 vector using forward primer 5’-TCGATGAATTCggtggcggtggctcgggcggaggtgggtcgATGGTGAGCAAGGGC -3’ and reverse primer 5’-TCGATGTCGACTTACTTGTACAGCTCGTC -3’, then ligated into pBabe-puro vector (Addgene #1764) between EcoRI and SalI sites. After the EcoRI site, the forward primer contained nucleotides encoding a 10-amino acid flexible linker sequence (GGGGSGGGGS) between the KIF18B and GFP sequences. KIF18B was amplified from pEGFP-KIF18B (Gift from Claire Walczak) using forward primer 5’-TCGATGGATCCGCCACCATGGCAGTGGAGGACAGC-3’ and reverse primer 5’-TCAGTGAATTCGTGCCAGTCACCTGG-3’ and inserted between BamHI and EcoRI sites. The stop codon was removed from the KIF18B C-terminal end. The primers were designed against Homo sapiens kinesin family member 18B (KIF18B), transcript variant 2, mRNA (NM_001264573.2) to produce the insert. Successful cloning was confirmed by DNA sequencing via Eton Biosciences (Research Triangle Park, NC).

Retrovirus was created using Phoenix cells, Mirus TransIT-LT1 reagent, and the provided protocol (Mirus Bio, MIR 2300). Virus was filtered with 0.45-μm filter before use. To establish a stably-expressing cell line, 1 x 3.5 cm plate of KIF18BKO cells were incubated with 2 mL KIF18B-GFP retrovirus, polybrene (6 μg/mL), and FBS (9%) for 15 min at room temperature. The plate was spun at 32 °C in a benchtop centrifuge for 30 min at 1,100 x g, rinsed 3 x 1X PBS, and then incubated with E low calcium medium + Dual SMAD inhibitors. Cells were passaged and expanded, then selected using puromycin. After selection, cells were sorted for GFP+ expressors.

### Immunoblots

Cells were lysed and collected on ice in RIPA Buffer (50 mM Tris pH 7.9, 150 mM NaCl, 0.5 mM EDTA pH 8.0, 1% NP-40, 0.1% SDS, 0.5% Sodium deoxycholate) with Protease Inhibitor Cocktail (Roche, 11697498001) and PMSF (1 mM, PanReacAppliChem, A0999). Samples were then rotated at 4 °C for 30 min, sonicated for 3 x 10 sec and placed on ice in between rotations. Protein was separated from cellular debris via centrifugation (10 min, 15,000 x g at 4 °C) and stored at −80 °C.

Lysates were mixed 1:1 with loading buffer (15% β-Mercaptoethanol, 10% SDS, 40% glycerol, and 3% Bromophenyl Blue) for protein gel analysis. Protein samples were then denatured by boiling for 5 min at 100 °C and subsequently run on 7.5% polyacrylamide gels at 100V for approximately 120 minutes. Proteins were then transferred from the gel onto nitrocellulose membranes at 100V for 80 minutes. Following transfer, membranes were block with 5% bovine serum albumin (BSA) in PBS-T (2 % Tween-20 in phosphate buffered saline) for 1 hr. Membranes were then incubated overnight in primary antibodies, rocking at 4 °C. Membranes were then washed 3 x 5 minutes in 1X PBS-T, incubated in secondary antibodies at room temp, rocking, for 1 hour, then washed 3 x 5 minutes in 1X PBS-T and visualized using a LI-COR Odyssey FC System. Primary antibodies (diluted in 5% BSA in 1X PBS-T) were as follows: Mouse anti-β-Actin (1:10,000, Sigma-Aldrich, A5441), Rat anti-α Tubulin (1:2000, Santa Cruz, sc-53029), Rabbit anti-KIF18B (1:500, Sigma-Aldrich, HPA024205), Chicken anti-GFP (1:5000, Abcam, ab13970). Secondary antibodies were Licor, IRDye 680RD Series or CW800 Series diluted 1:5000 in 5% BSA/1X PBS-T. For GFP antibody visualization, membrane was incubated with secondary Goat anti-Chicken HRP (1:5000 in 5%BSA/1X PBS-T, Jackson ImmunoResearch, 103-035-155) then incubated for 2 min in the dark with ECL Western Blotting Substrate (ThermoFisher, PI-32106) before using the LI-COR system. The LI-COR system was used for quantifications. β-Actin was used as a loading control and compared to KIF18B for protein-fold quantification.

### Immunofluorescence

#### Tissue Culture, Sample Collection through Primary Antibody

Cells were cultured on glass coverslips. For assays using spindle orientation machinery, cells were plated on laminin-coated coverslips (100 μM, Invitrogen, 23017015). Coverslips were quickly rinsed in warm 1X PBS, then fixed in ice-cold methanol (MeOH) for 3 min. MeOH-fixed coverslips were then rinsed with 1X PBS-T (0.2% Triton-X) for 5 min. For fixation of microtubules, coverslips were fixed in 37 °C glutaraldehyde fixative (80 mM PIPES pH 6.9, 50 mM NaCl, 2 mM MgCl2, 0.2 % Triton-X, 1% Glutaraldehyde) for 10 min followed by rinsing with 1X PBS-T 3 x 5 min. Glutaraldehyde was then quenched with a small amount of sodium borohydride in PBS for 7 min followed by 3 x 5 min rinse in 1X PBS-T.

After rinses, cells were blocked with “Block” (3% BSA/5% Normal Donkey Serum (NDS)/5% Normal Goat Serum (NGS) in PBS-T) or “NDS Block” (3% BSA/5% NDS in PBS-T) for 15 min. All antibodies were diluted in blocking solution and incubated at conditions indicated in the table below. Following primary incubation, coverslips were rinsed 3 x 5 min in PBS-T.

#### Mouse Tissue, Tissue Collection through Primary Antibody Incubation

Tissue was embedded in OCT (Sakura Finetek, 4583), frozen, and sectioned at 10 μm using a cryostat. Sections were fixed in 4% PFA for 8 min at RT followed by a 5 min wash in 1X PBS-T and blocking in “Block” or “NDS Block” for 15 min (See above paragraph). All antibodies were diluted in blocking solution and incubated at conditions indicated Table 1. Following primary antibody incubation, sections were rinsed 3 x 5 min in 1X PBS-T.

**Table 1.** Primary Antibody and Immunofluorescent Staining Information.

| Antibody | Host | Supplier | Catalog Number | Fixative | Dilution | Incubation & Notes |
| --- | --- | --- | --- | --- | --- | --- |
| β-4 Integrin | Rat | BD Biosciences | 553745 | PFA | 1:200 | 1 hr, RT |
| α-tubulin | Rat | Santa Cruz | sc-53029 | MeOH/Glut | 1:100 | 1 hr, RT |
| Foxi3 (L-17) | Goat | Santa Cruz | sc-324865 | PFA | 1:100 | 1 hr, RT |
| GFP | Chicken | Abcam | ab13970 | MeOH/PFA | 1:200 | 1 hr or O/N, 4 °C |
| KIF18B | Rabbit | Sigma | HPA024205 | MeOH | 1:100 | O/N, 4 °C. Pre-extract in 0.5% Triton-X for 30s at 37 °C. |
| KIF18B | Gift | Medema Lab,<br>Netherlands Cancer Institute | N/A | PFA | 1:50 | O/N, 4 °C |
| LHX2 (C-20) | Goat | Santa Cruz | sc-19344 | PFA | 1:100 | 1 hr, RT |
| NuMA | Rabbit | Thermo Fisher | PA3-16829 | MeOH | 1:100 | O/N, 4 °C |
| NuMA | Goat | Santa Cruz | sc-51164 | MeOH | 1:100 | O/N, 4 °C |
| p150<sup>glued</sup> | Mouse | BD Biosciences | 610473 | MeOH | 1:100 | O/N, 4 °C |
| Sox2 | Rabbit | Abcam | ab92494 | PFA | 1:200 | 1 hr, RT |
| Sox2 | Rat | Invitrogen | 14-9811-80 | PFA | 1:200 | 1 hr, RT |
| Sox9 | Rabbit | Millipore | Ab5535 | PFA | 1:1000 | 1 hr, RT |

#### Tissue Culture and Mouse Tissue, Secondary Antibody though Mounting

All samples were incubated in secondary antibodies (Table 2) diluted in Block/NDS Block plus Hoechst (Invitrogen, H3570) for 15 min at room temperature. Samples were then washed 3 x 5 min in PBS-T. Coverslips were mounted with antifade solution (0.25% *p*-phenylenediamine, 10% 1X PBS, 90% glycerol, pH 9.0) and sealed with clear nail polish.

**Table 2.** Secondary Antibody Information.

| Antibody | Supplier | Catalog Number | Dilution |
| --- | --- | --- | --- |
| Alexa Fluor 488 Donkey anti-Chicken | Jackson ImmunoResearch | 703-545-155 | 1:200 |
| Alexa Fluor 488 Donkey anti-Goat |  | 705-545-003 |  |
| Alexa Fluor 488 Donkey anti-Rabbit |  | 711-545-152 |  |
| Alexa Fluor 488 Donkey anti-Rat |  | 712-545-153 |  |
| Alexa Fluor 647 Donkey anti-Rabbit |  | 711-605-152 |  |
| Alexa Fluor 647 Donkey anti-Rat |  | 712-605-153 |  |
| Rhodamine Red- X Donkey Anti-Goat |  | 705-295-147 |  |
| Rhodamine Red- X Donkey Anti-Mouse |  | 715-295-151 |  |
| Rhodamine Red- X Donkey Anti-Rabbit |  | 711-295-152 |  |
| Rhodamine Red- X Donkey Anti-Rat |  | 712-295-153 |  |

### Spindle Orientation Analysis

For spindle orientation analysis of fixed cells, cells were plated on laminin-coated coverslips. 0.5 mM calcium was added 4 hours prior to fixation to upregulate cellular junctions and induce proliferation. FIJI (ImageJ) was used to process images. Spindle angles were measured using max-intensity projections of all z-slices that contained cortical spindle orientation machinery. Division angles were measured as the angle between spindle pole NuMA staining and the center of the cortical NuMA crescent. Only metaphase cells were used for analysis.

For spindle orientation analysis of back skin tissue sections, division angles of basal cells were measured between NuMA-GFP puncta with respect to the basement membrane, marked by β-4 integrin. Placodes cells were identified by morphological characteristics and by Sox2 to mark the dermal condensate (Saxena et al., 2019). Both metaphase and anaphase cells were used, as identified by their DNA condensation.

### Time-lapse Imaging

KIF18B;EB1-GFP control and KO primary keratinocytes were isolated from mouse back skin as described above and were plated on 3.5 mm glass-bottomed TC plates (MatTek Corporation, P35G-1.5-14-C) coated with laminin. 0.5 mM calcium was added 4 hours ahead of imaging to upregulate cellular junctions and induce proliferation. Cells were imaged on an Andor XD Revolution Spinning Disk Confocal with temperature control chamber (37 °C) and CO_2_ (5%) (Duke Light Microscopy Core Facility). 30 sec movies (1 frame/second) were taken using 60x Plan-Apo 1.2 NA water objective, Andor Ixon3 897 512 EMCCD camera and MetaMorph Software. Post-acquisition processing of movies was done with FIJI (ImageJ). EB1-GFP comets were tracked manually using FIJI Manual Tracking Plugin. 10 EB1-GFP puncta were tracked per cell, 5 per spindle pole but across the entire astral region. The brightest puncta were selected for tracking.

### Microscopy

All fluorescent images of fixed samples were taken using a Zeiss AxioImager Z1 microscope with an Apotome 2 attachment, PlanAPOCHROMAT 20X objective, 40X/1.3 oil objective, or Plan-NEOFLUAR 63X/1.4 oil objective and Axiocam 506 monocamera and Zeiss Zen Software. Images were processed using FIJI (ImageJ) and Adobe Photoshop.

### Image Quantification, Graphing and Statistical analysis

Image quantifications were done using either LI-COR Odyssey FC System (western blot) or FIJI. All graphing and statistical analyses were done using Prism, except for spindle orientation analysis. Radial histograms for spindle orientation were done using MATLAB and analyzed for significance with a Kolmogorov-Smirnov test. All remaining data was statistically analyzed using a Student’s unpaired t-test.

## Acknowledgments

We would like to thank Claire Walczak (Indiana University), René Medema (Netherlands Cancer Institute), and Purushothama Rao Tata (Duke University) for reagents. We thank Julie Underwood for care of the mice and the entire Lechler lab for their thoughtful comments throughout the course of this project and on the manuscript.

---

**Supplemental Figure 1.**
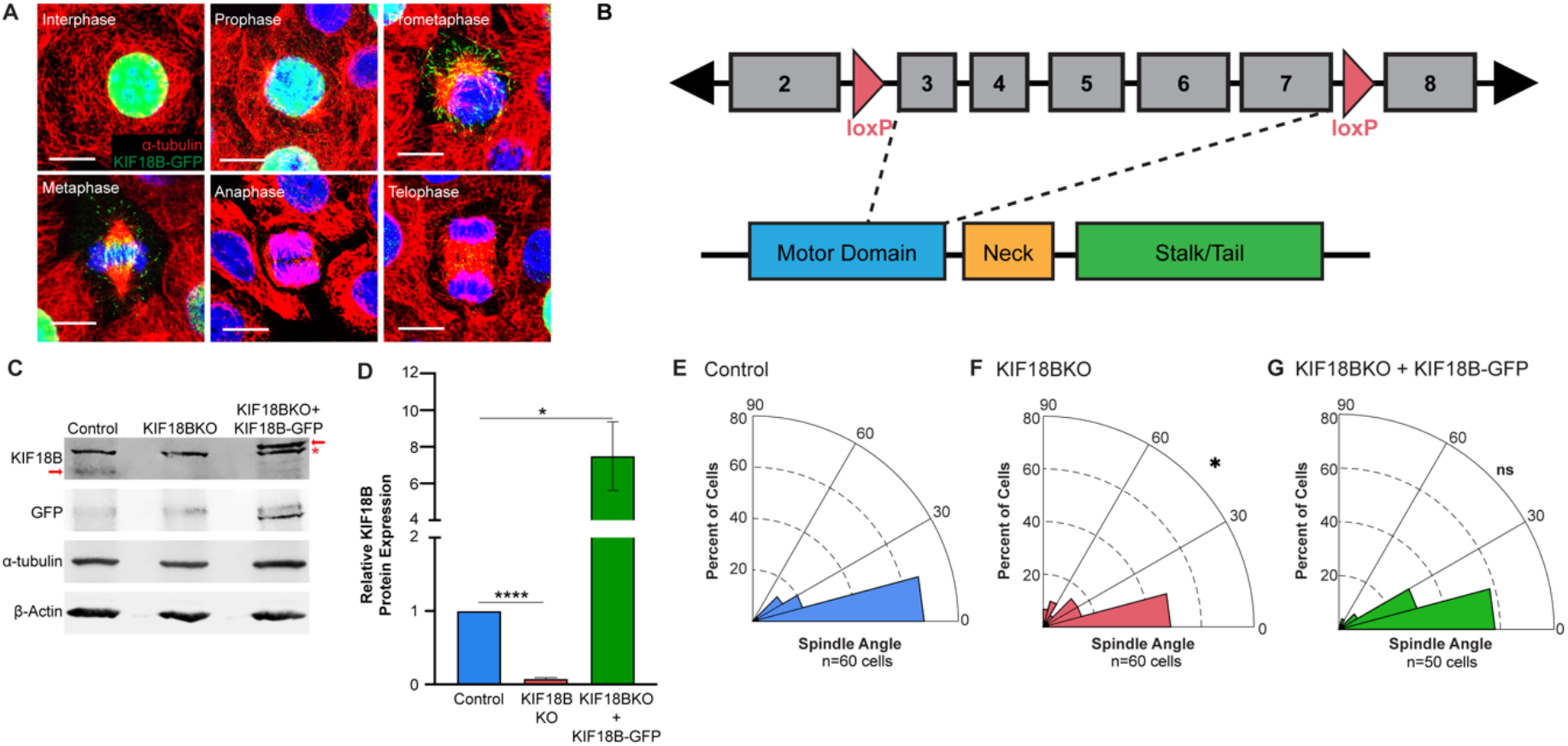
KIF18B is required for spindle orientation in an additional KIF18BKO line. Related to Figure 1. (A) KIF18B-GFP (green) localization pattern through the cell cycle. α-tubulin (red) labels microtubules. (B) Diagram of gene-specific portion of targeting vector for Kif18b conditional knockout. (C) Western blot of proteins prepared from cell lysates for control, KIF18BKO, and KIF18B-GFP mouse keratinocyte cell lines (Cell Line 2). Cell Line 2 was generated independently from the first set of control and KIF18B knockout cells. Red arrows indicate Kif18b, while the asterisk denotes a non-specific band. (D) Quantification of KIF18B protein levels in cell lines (n=3, Control vs KIF18BKO p<0.0001, Control vs. KIF18B-GFP p=0.0055, Student’s unpaired t-test). (E-G) Radial histograms of spindle orientation in control (E), KIF18BKO (F), and KIF18B-GFP cells (G) (Control vs. KIF18BKO p = 0.047, Control vs. KIF18B-GFP p = .9479, Kolmogorov-Smirnov Test). Scale bar = 10 μm.

**Supplemental Figure 2.**
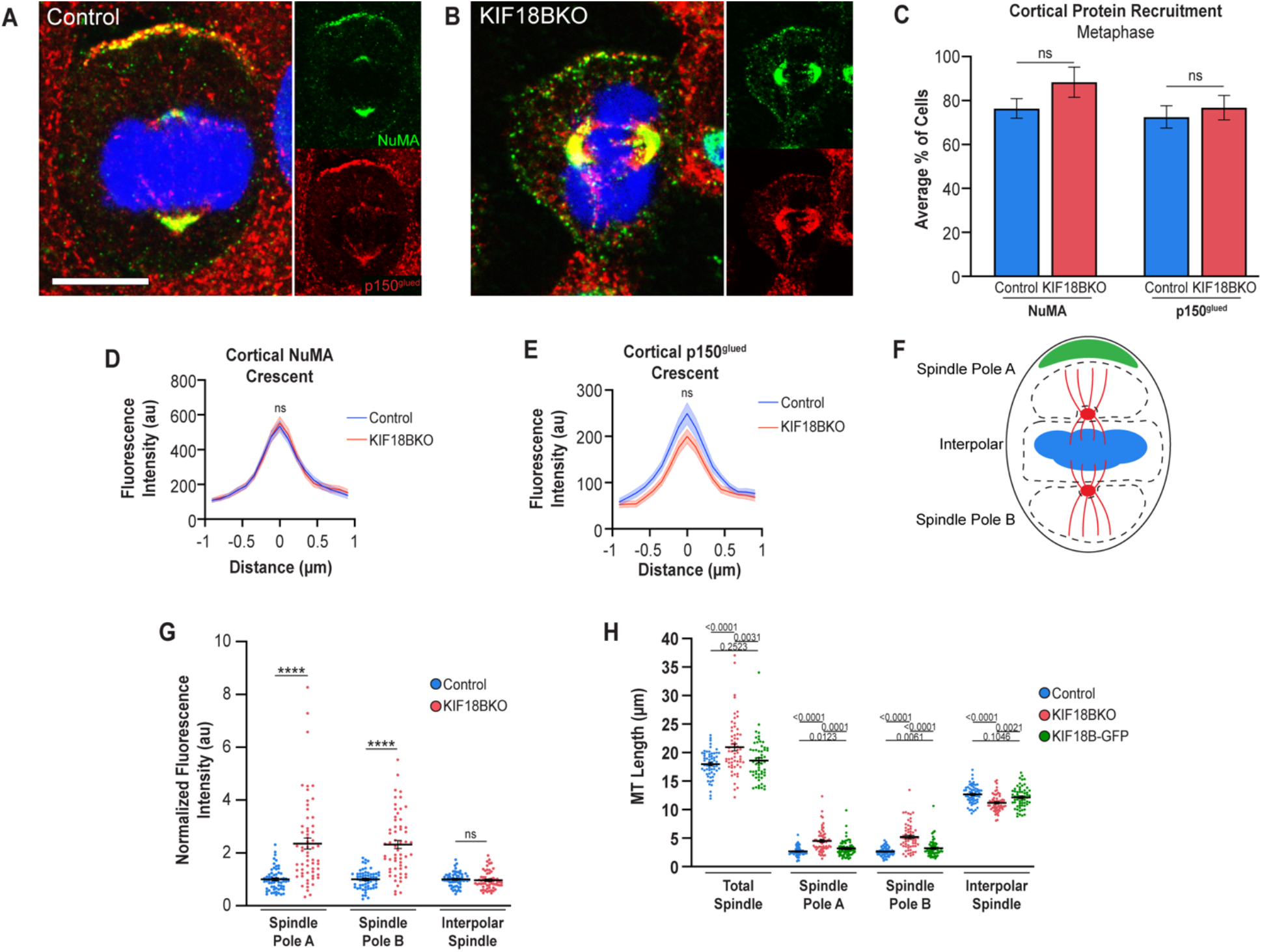
KIF18B does not affect spindle orientation machinery recruitment. Related to Figure 2. (A-B) Example images of NuMA (green) and p150glued (red) cortical protein recruitment in control and KIF18BKO cells. (C) Quantification of percent average of cortical protein recruitment in metaphase cells. (n= 56 control cells, 53 KIF18BKO cells, NuMA p= 0.9533, p150glued p=0.1857). (D-E) Fluorescence intensity measurements of cortical protein across center of protein crescent. (n=30 cells/condition, p = ns). (F) Example diagram of measurements for fluorescence intensity and microtubule length. Microtubules are labeled in red. Centrosomes were left out of quantifications due to overexposed intensity. Spindle pole A denotes side of spindle pole with cortical NuMA crescent. (G) Quantification of microtubule fluorescence intensity at indicated spindle location, normalized to control cell average (n= 60 cells/condition, Astral Microtubules p<0.0001, Interpolar Spindle p=0.5614, Student’s unpaired t-test). (H) Quantification of spindle length at indicated location in control, KIF18BKO, and KIF18BKO + KIF18B-GFP cells (n=60 cells/condition, Total Spindle p<0.0001, Astral Microtubules p<0.0001, Interpolar Spindle p<0.0001, Student’s unpaired t-test).

**Supplemental Figure 3.**
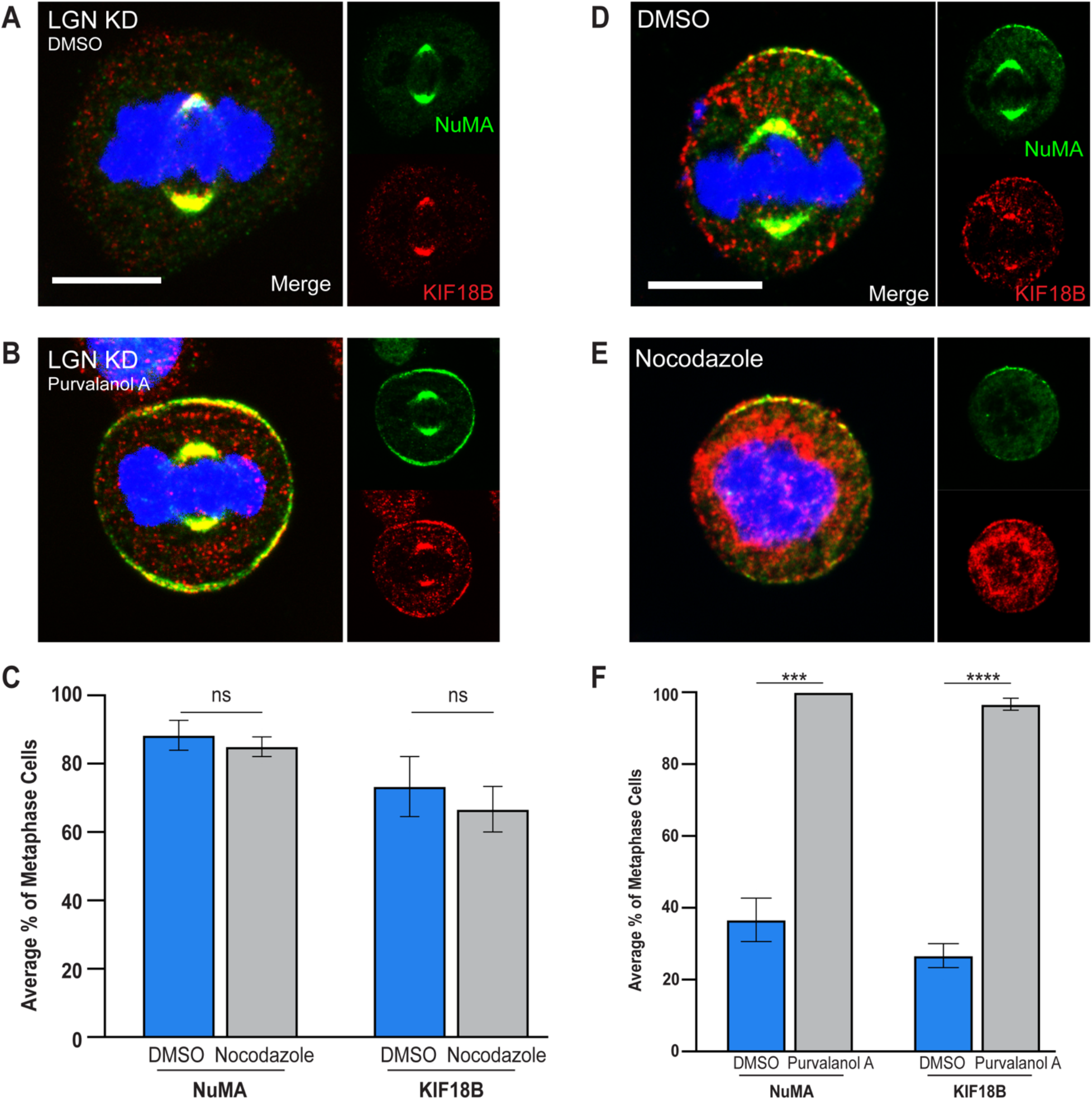
KIF18B localization is similar to that of NuMA. Related to Figure 5. (A-B) Example images of NuMA (green) and KIF18B (red) cortical protein recruitment in LGN KD cells during metaphase in control or Purvalanol A treated cells. (C) Quantification of cortical protein recruitment (n=60 cells/condition, NuMA p=0.0005, KIF18B p<0.0001, Student’s unpaired t-test). (D-E) NuMA (green) and KIF18B (red) localization at the cell cortex with DMSO (D) or nocodazole treatment (E) to depolymerize microtubules. (F) Quantification of average percent of cortical protein recruitment during metaphase in control or treated cells. (n=60 cells/condition, NuMA p=0.5614, KIF18B p=0.5790, Student’s unpaired t-test). Scale bars = 10 μm.

**Supplemental Figure 4.**
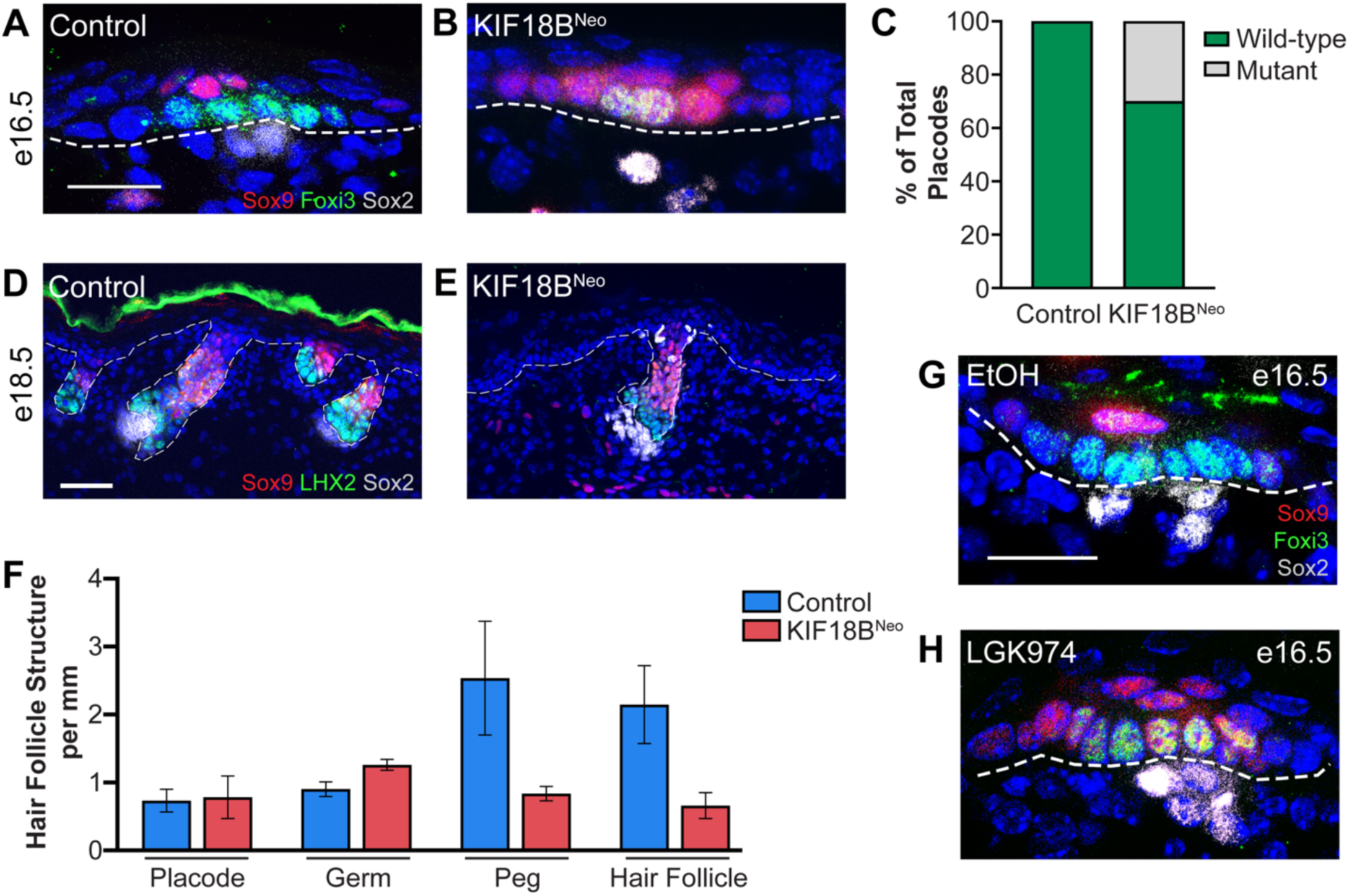
KIF18B mutant has reduced number of hair follicles. Related to Figure 6. (A-B) Images of control and KIF18B mutant placodes. Foxi3 (green) marks basal placode cells and Sox9 marks progenitor cells. Scale bar is 20 μm (C) Quantitation of percent of mutant placodes in KIF18B^Neo^ mutant versus control at e16.5. (n=20 placodes). (D-E) Back skin of control and KIF18B^Neo^ mutant back skin at e18.5. Sox9 (red) marks progenitor cells, LHX2 marks hair follicle cells, and Sox2 marks the dermal papillae/condensates. Scale bar is 50 μm. (F) Quantitation of number of hair follicle structures in KIF18B^Neo^ mutant compared to control (n=2 mice per condition). (G) Example placode of mouse treated with DMSO. Foxi3 (green) marks basal placode cells, Sox9 marks progenitor cells, and Sox2 marks dermal condensate. Scale bar is 20 μm. (H) Placode of mouse treated with single intraperitoneal injection of Wnt inhibitor LGK974. In all images above, dotted line marks basement membrane.

